# Snake Herders: Novel anti-predator behavior by black-tailed prairie dogs in response to prairie rattlesnakes

**DOI:** 10.1101/2022.07.16.500317

**Authors:** J.L. Verdolin, Ellen K. Bledsoe

**Affiliations:** School of Natural Resources and the Environment, University of Arizona, Tucson AZ, USA; Department of Biology, Duke University, Durham, NC, USA

**Keywords:** anti-predator behavior, prairie dog, behavior, predator-prey interaction, Cynomys

## Abstract

We describe a case of a unique antipredator behavior sequence in response to rattlesnakes observed in a population of black-tailed prairie dogs in Fort Collins, Colorado. An analysis of 5 independent opportunistic video recordings of 6 identifiable individuals revealed individuals across multiple social groups within the population engaged in novel behavioral responses to prairie rattlesnakes. Using Markov-chain analysis we found that prairie dogs engaged in non-random behavioral transitions and that specific pairs of behaviors were contributing to this pattern. We also observed that prairie dogs in this population engaged in novel responses to prairie rattlesnakes, including ‘escorting’ behavior, where a prairie dog would walk alongside the snake.

## Introduction

Predation is a significant selective force exerting pressure on prey populations to evolve counter-measures (Edmunds 1974; Endler 1986). Within a species there can be considerable variation in antipredator strategies reflecting adaptation to local predator communities (Urban et al. 2020). The development and persistence of behavioral defenses is expected to balance costs and benefits, such that excessively costly behaviors will not persist (Cooper and Frederick, 2010). In general, optimal escape theory would predict that the benefit of confronting a lethal predator must outweigh the potential loss of fitness, in this case death (Ydenberg and Dill, 1986).

Black-tailed prairie dogs (*Cynomys ludovicianus*) are found from Chihuahua in northern Mexico to Saskatchewan, Canada, and occupy grassland environments (Slobodchikoff et al. 2009a). Like other species of prairie dog, black-tails are predated on by a suite of predators. Consequently, prairie dogs as a group are an excellent system to study the development and persistence of antipredator behaviors. Past research has documented complex vocal and behavioral responses in prairie dogs that reduce predation risk (Slobodchikoff et al. 1991; Slobodchikoff 2002; Slobodchikoff and Placer 2006; Frederickson and Slobodchikoff 2007; Slobodchikoff et al. 2009b). For example, the alarm calls of Gunnison’s prairie dogs (*Cynomys gunnisoni*) differ depending on the type of predator and the speed of approach (Kiriazis, 1991; Slobodchikioff et al. 1991; Slobodchikoff, 2002). In addition, behavioral escape strategies in prairie dogs correspond to the alarm call emitted in response to different types of predator and juveniles learn through exposure to adults in the community (Kiriazis and Slobodchikoff 2006; Shier and Owing 2007; Frederikson and Slobodchikoff 2010).

In areas where prairie dogs are subjected to similar predator communities, it might be anticipated that their antipredator strategies will be comparable. Across populations, however, we find evidence for variability in both the vocal and behavioral responses exhibited in response to predators. For instance, there is strong evidence for geographic variability of prairie dog alarm calls at local and regional scales (Slobodchikoff et al. 1998; Perla and Slobodchikoff 2002). More recently, Dennis et al. (2020) reported that black-tailed prairie dog colonies more than 30 km from each other exhibit distinct alarm call dialects.

Variability in behavioral engagement with predators has also been documented, particularly with snakes. In some populations prairie dogs attacked nonvenomous bullsnakes (*Pituophis catenifer sayi*) more frequently than rattlesnakes (*Crotalus spp*.) and spent more time in snake-directed behaviors toward rattlesnakes, while in another population prairie dogs behave similarly toward both bullsnakes and rattlesnakes (Owings and Loughry 1985; Loughry 1988, 1989). Loughry (1989) concluded that the degree of exposure and experience a colony has with snakes contributes to this difference. This is also similar to what has been observed in other species of ground squirrels (Owings et al. 2001). Yet, unlike other species of ground squirrels (e.g., rock squirrels (*Otospermophilus variegatus*) and California ground squirrels (*Otospermophilus beecheyi*)), there is no evidence that adult prairie dogs are immune to the venom of prairie rattlesnakes (Owings 2001), making direct confrontation with a rattlesnake potentially deadly.

Here, we report on a distinct combination of anti-predator behaviors in response to prairie rattlesnakes (*Crotalus viridis*) at Coyote Ridge Natural Area in Ft. Collins, Colorado. We explored whether there was a non-random sequence to the behaviors and, if so, which behavior transitions were responsible for driving the non-random pattern.

## Methods

### Data Collection

JLV made direct opportunistic observations from July 9, 2021 to July 13, 2021, sitting in the open alongside the Coyote Ridge Trail at Coyote Ridge Natural Area. JLV recorded all observations with a Canon PowerShot SX70 HS Digital Camera. Six different individual prairie dogs were recorded interacting with and responding to prairie rattlesnakes in the colony. Individual prairie dogs were identified with unique marks, such as scars, bald spots, and other distinguishing features.

For each video, the individual, sex, date, time, bout, and video sequence was recorded by JLV to distinguish between different prairie dogs and interactions with rattlesnakes. For each bout, behaviors and behavioral transitions were recorded only when individuals were fully visible in the field of view.

### Data Analysis

All statistical analyses were conducted in R 4.1.2 (R Core Team, 2020). The data and R scripts for all analyses can be found on GitHub (https://github.com/bleds22e/SnakeHerders). We created a behavior transition matrix for prairie dogs, tabulating all the instances in which one behavior led to a different behavior. We excluded behaviors if 1) the behavior was not related to the prairie dog-snake interaction (e.g., greet-kiss or foraging), and/or 2) the behavior only occurred only once (e.g., fake-foraging). We used single-order Markov chains to test whether the behavioral transitions were non-random using the ‘verifyMarkovProperty’ function from the package ‘markovchain’ (Spedicato, 2017).

We were also interested in evaluating which transitions contributed to the nonrandom pattern. To do this, we ran chi-squared tests on the transition matrices to determine overall significance. Using the ‘FTres’ function from the ‘kequate’ package (Andersson et al. 2013), we used the values from the transition matrix and expected values calculated for the chi-squared test to determine the Freeman-Tukey residuals for each transition (Chen et al. 2002). From there, we calculated correlation values and significance values for each transition probability using the Freeman-Tukey residual values (’Hmisc’ package; Harrell 2022).

## Results and Discussion

### Prairie Dog Anti-snake Behavior

We observed a total of 6 interactions between prairie dogs and snakes. In our preliminary review of the video recordings, we noted several novel behaviors which have not previously been reported in the literature in response to predators, including snakes. Specifically, we observed prairie dogs engaged in foreleg drumming, back leg scratching, and “escorting”, where a prairie dog would walk alongside the rattlesnake for some period of time. These behaviors appeared to occur in a sequence and after a prairie dog initially approached the rattlesnake. For example, an individual approached the rattlesnake and patted the ground with her front feet (foreleg drumming), then she rocked and sniffed simultaneously, and then scratched herself with her back right leg (Figure 1A-C). Once the snake began moving, she walked alongside the snake, escorting the snake (Figure 1D).

**Figure 1.**
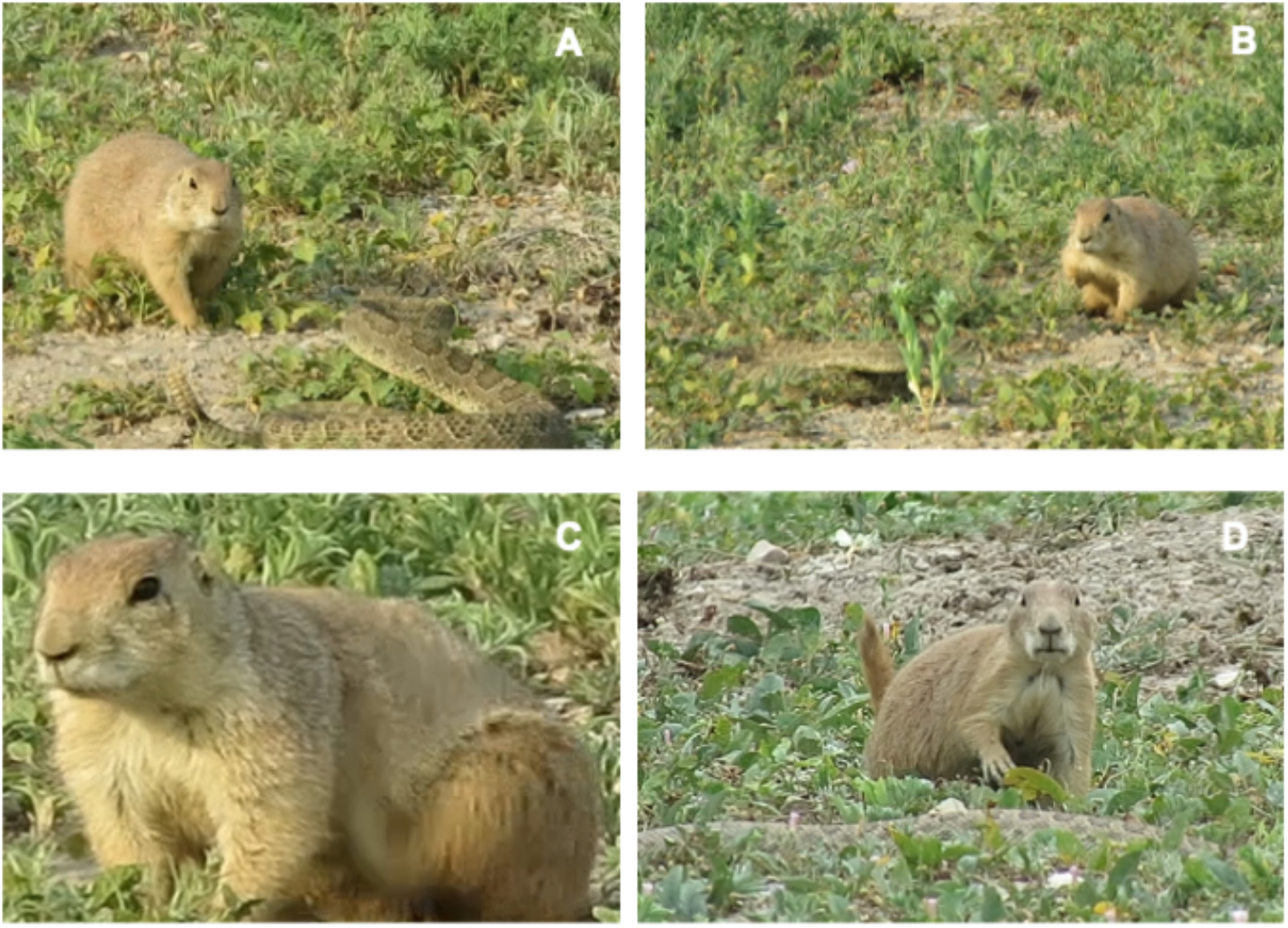
**A-D**.Observed behavioral sequence. Behavioral sequence of an interaction between a prairie dog and a rattlesnake. In this sequence, a female engaged in foreleg drumming (A), rocking and sniffing (B), and then back leg scratching (C). The female begins escorting the rattlesnake (D).

In addition to these novel behaviors, we also observed behaviors that have been previously described in response to predators such as rattlesnakes and bullsnakes. For example, we observed prairie dogs jump-yipping in proximity to the snake’s head as reported by Loughry (1989).

The Markov chain analysis of prairie dog behavioral transitions indicated a non-random pattern of behavioral sequences (*X*^2^ = 75.27, df = 36, *p* = 0.00014). The Pearson’s Chi-squared test was also significant (*X*^2^ = 353.6, df = 121, *p* < 0.0001), meaning the number of behavior transitions differed from expected. Subsequent pairwise comparisons using Spearman’s rank correlation revealed the specific transitions that were contributing to these results (Figure 2; Table 1). Notably, there were four significant behavioral transitions: escorting → retreat (r_s_ = 0.64, *p* = 0.024); jump-yip → retreat (r_s_ = -0.65, *p* = 0.022); rocking → rocking and sniffing (r_s_ = 0.71, *p* = 0.010); and foreleg drumming → jump back (r_s_ = 0.66, *p* = 0.020). Four additional behavioral transitions were borderline significant: rocking and sniffing → scratching (r_s_ = -0.54, *p* = 0.071); approach → jump-yip (r_s_ = 0.52, *p* = 0.085); jump-yip → rocking (r_s_ = 0.51, *p* = 0.090); and escorting → foreleg drumming (r_s_ = 0.51, *p* = 0.090).

**Figure 2.**
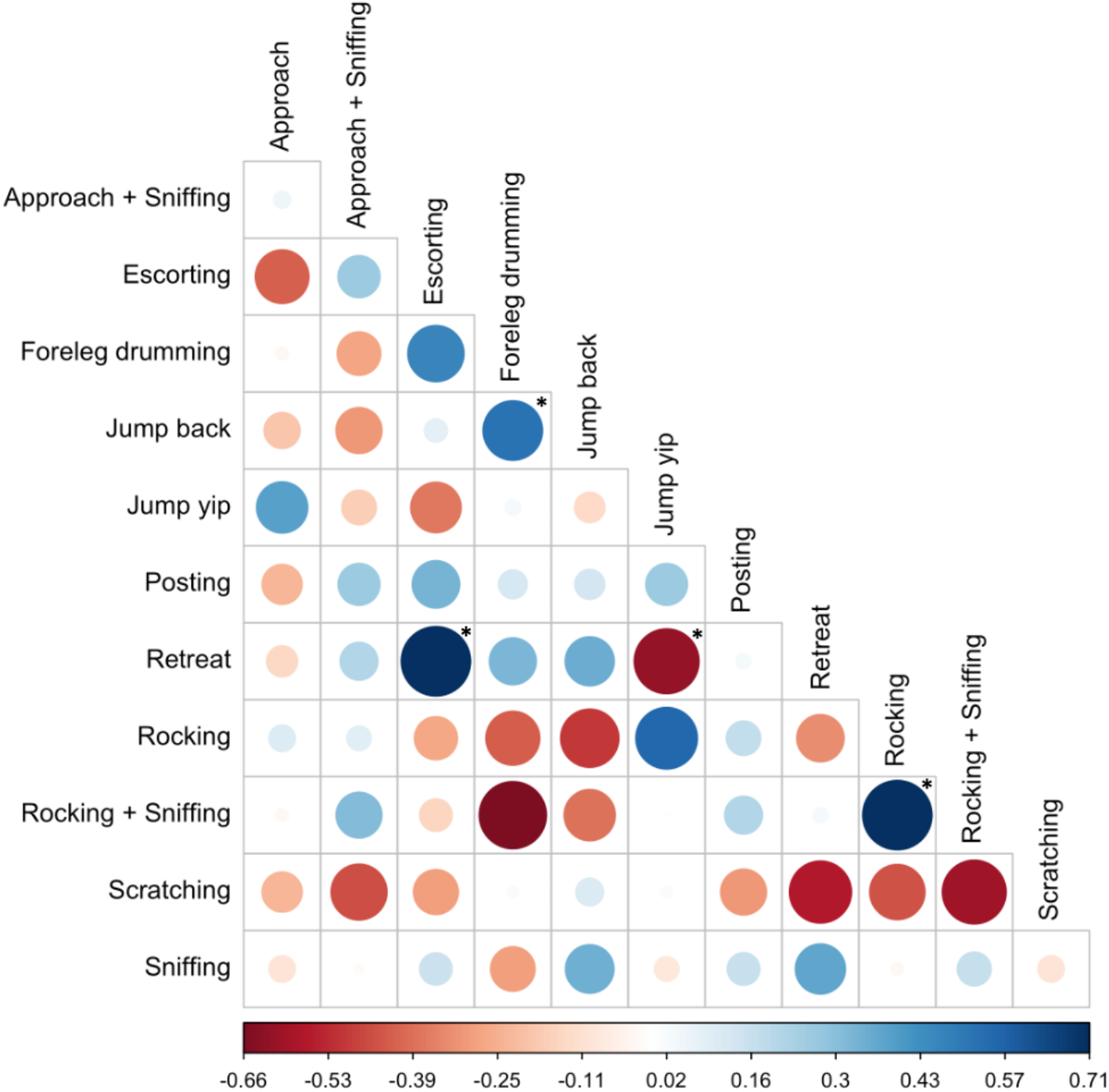
Freeman-Tukey pairwise comparison. Freeman-Tukey residuals for each behavioral transition pair. Transitions are from the behavior on the x-axis to the behavior on the y-axis; for example, the transition state of the box in the upper left-hand corner is an “approach” to “approach and sniff.” Color of dots indicates the value of the Freeman-Tukey residuals for each transition pair, calculated from the observed and expected values of the transition matrix. Positive (blue) values indicate that the transition occurred more frequently than would be expected, while negative (red) values indicate the transition occurred less frequently than expected. Size of dots corresponds to the absolute values of the Freeman-Turkey residuals. Asterisks indicate significant transition pairs.

When we looked at the Freeman-Tukey residuals for each significant transition pair, all except jump-yip → retreat occurred more frequently than expected (positive r_s_ values). Together, these behavioral transitions appear to be driving the non-random pattern detected by the Markov-Chain analysis but in different ways. For example, escorting transitions to retreating more frequently than expected, while retreating happens less frequently than expected after jumps-yipping (Figure 2). The reason for this may be because while a prairie dog is escorting, the snake is moving out of the area making it safe to retreat. However, when jump-yipping, the prairie dog is still engaged in a display toward the rattlesnake, thus is less likely to retreat if the rattlesnake is still present.

Although the prairie dogs are engaged in non-random behaviors in response to rattlesnakes, and we have identified specific behavioral transitions that occur more or less often than expected, we recognize we have a small sample size involving six individual prairie dogs and 189 total behavioral transitions.

### Prairie Dog Snake Detection

In several instances it was clear the prairie dogs were trying to determine the exact location of the snake. This was done by sniffing, rocking and sniffing, or elongating the head and neck while moving it up and down. Although prairie dogs are known to have color vision, it is difficult for them to discriminate in the yellow and green spectrum (Cain and Carlson 1968; Slobodchikoff et al. 2009b). Prairie rattlesnakes are well camouflaged and, when present in grass, may be especially difficult for prairie dogs to see until they are very close to the snake.

We suspect that prairie dogs may be using olfaction to identify the specific location of a rattlesnake before engaging in additional snake-directed behaviors. We observed prairie dogs rocking and sniffing, cautiously approaching the snake, and then quickly jumping back as soon as they were in close proximity (approximately <15 cm), after which they approached the snake more confidently. Once the snake’s location was determined, prairie dogs remained in this range of proximity during the majority of the interaction.

On one occasion there was a clear flehmen response by a focal prairie dog, further pointing to chemosensory information being used to locate the snake. Most organisms, including snakes, have a natural scent and there is evidence that other species can detect the odor of rattlesnakes. For example, California ground squirrels have been observed actively applying rattlesnake scent from shed snakeskin to their fur (Clucas et al. 2008), a behavior not observed in this or other populations of prairie dogs. Thus, it seems likely that prairie dogs were cueing in on the natural scent of the rattlesnake as part of their antipredator strategy.

### Prairie Dog Manipulation of Prairie Rattlesnakes

We discovered that when a focal prairie dog engaged in foreleg drumming, scratching, or jump yipping, the snake responded by moving its body, orienting its head, or raising its head in the direction of the prairie dog. Unfortunately, there were not enough behavioral transitions to analyze the prairie dog-snake interaction data quantitatively. However, it appears that the behavior of the prairie dogs elicited a response from the snake to orient and begin moving toward the prairie dog. Once the snake was moving, the focal prairie dog shifted their position, walking alongside, or escorting, the snake. This escorting behavior has not been reported in other prairie dog populations and may represent a novel anti-predator behavior unique to this colony.

Prairie dogs surrounding the focal prairie dog did not alarm call or assist the individual in any way. In three instances, a second prairie dog was in close proximity (<0.5 meter) and was either observing, posting, or foraging. This is markedly different from reports of multiple prairie dogs attacking and mobbing both rattlesnakes and bullsnakes (Tang and Haplin 1983). On two occasions an adult female had juveniles in close proximity that were observing the interaction. Subsequently, they engaged in rocking and sniffing behavior. Given the close proximity of other adults and occasionally juveniles, there is an opportunity for learning through observation and exposure. This is similar to reports by Owings and Loughry (1985) that prairie dogs nearby monitor the snake-directed behavior of others.

Also of note is that the focal individual was consistently in close proximity (<15 cm) to the rattlesnake during the majority of the interaction. We do not think prairie dogs in this population remain close in order to attack the snake, as there were only two instances where a prairie dog struck the snake (Figure 3D). However, this distance is clearly within striking distance for prairie rattlesnakes. This makes proximity to, and direct interaction with, rattlesnakes potentially deadly. On July 9th, JLV observed a swollen, bluish-colored deceased prairie dog on the outside of a burrow and another prairie dog rocking and sniffing at the lip of the burrow (Fig 3-AB). This is consistent with a prairie dog being exposed to rattlesnake venom. Several hours later a rattlesnake was observed attempting to consume the deceased prairie dog (Figure 3C). It remains unclear why adult prairie dogs engage in such high risk anti-predator behavior. One possibility is that adults are protecting smaller, more vulnerable pups. However we might expect that adults employ a different tactic outside of the reproductive season and early juvenile emergence period. To properly assess this we will focus future efforts on exploring whether or not there are seasonal differences in how prairie dogs respond to the presence of rattlesnakes. In addition, the close proximity of juveniles and other adults observing focal prairie dogs interacting with rattlesnakes hints at the possibility that the specific behavioral sequences observed in this colony are learned and socially transmitted.

**Figure 3.**
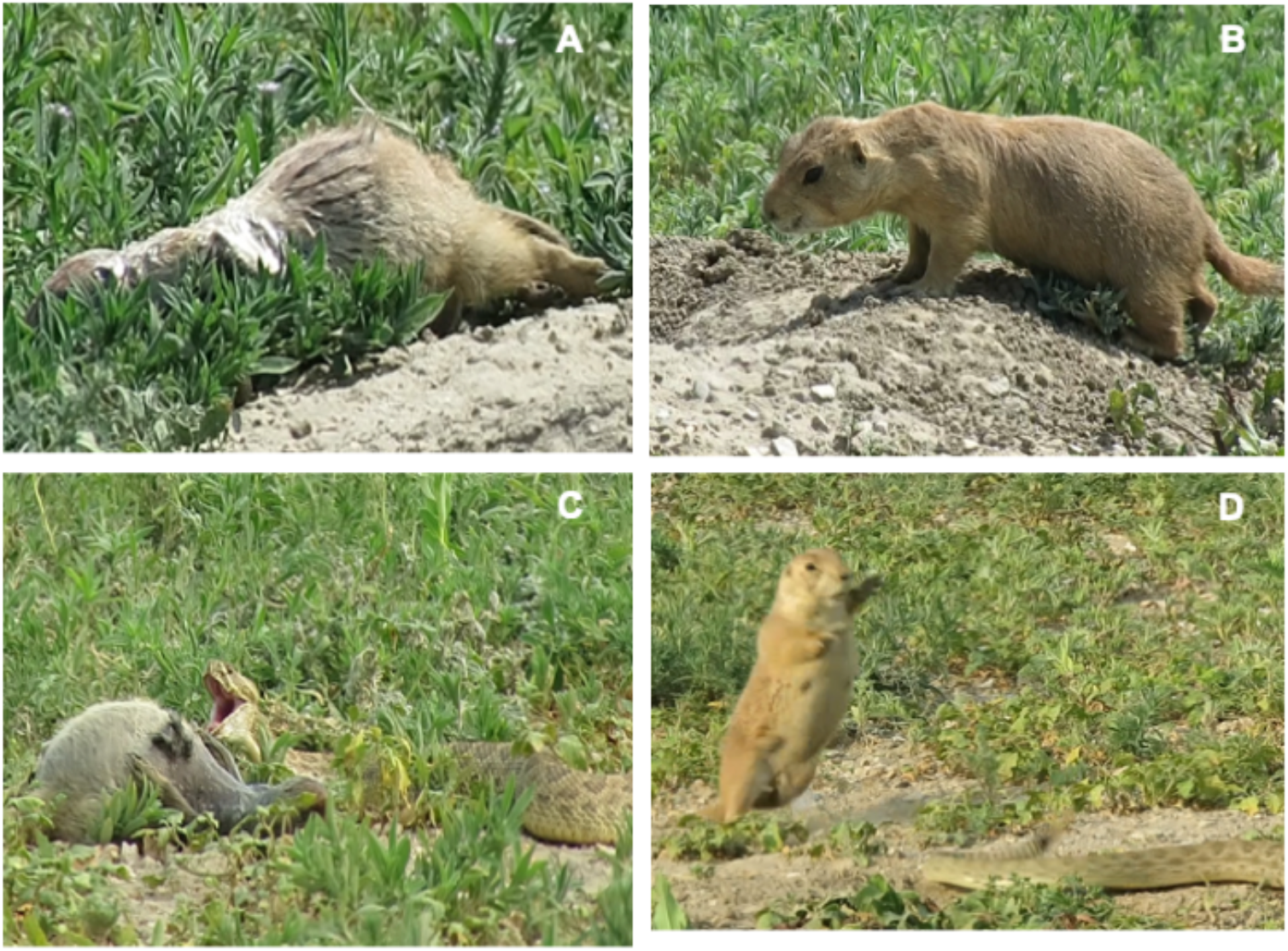
**A-D**. Prairie dogs engage in risky interactions with rattlesnakes. Panel A shows a deceased prairie dog on the edge of a burrow. Panel B is a second prairie dog investigating, sniffing, and rocking at the lip of the same burrow. Panel C shows the rattlesnake attempting to consume the deceased prairie dog. Panel D shows one occasion where a prairie dog struck the tail of the rattlesnake.

## Conclusions

We found that black-tailed prairie dogs on this colony engaged in anti-predator behavior in response to prairie rattlesnakes that included a combination of novel and previously documented behaviors. Specifically, we found that these prairie dogs exhibited novel non-random behavioral sequences that included foreleg drumming, scratching, and escorting.

Previous reports on interactions between black-tailed prairie dogs and rattlesnakes describe mobbing, attacks, and killing of snakes by prairie dogs (Tang and Haplin 1983), something not observed in this population. Despite exposing themselves to considerable risk, adult prairie dogs on this colony appear to use a set of structured behaviors to deal with a prairie rattlesnake in their territory. Most of the interactions were deliberate, appearing aimed at signaling to the snake, and encouraging and escorting the snake out of the focal prairie dog’s territory. Future studies will be aimed at exploring seasonal patterns to assess whether prairie dogs adjust their anti-predator strategies according to the presence of more vulnerable pups. Finally, we will quantify snake-directed behavioral variability across colonies and species of prairie dogs and determine if these behaviors are learned and socially transmitted within colonies.

